# Molecular determinants of differential substrate selection between the Src family kinases Lck and Src

**DOI:** 10.64898/2026.06.29.735195

**Authors:** Konstantina Karpouzou, Marco D’Abramo, Alessandro Grottesi, Oreste Acuto, Konstantina Nika

**Affiliations:** Department of Biochemistry, School of Medicine. University of Patras, Greece; Department of Chemistry, University of Rome “La Sapienza”, p.le A. Moro, 5, 00185, Rome, Italy; CINECA - Italian Computing Centre. SuperComputing Applications and Innovation Department (SCAI). Via dei Tizii 6 - 00185 Rome. Italy; Cell Signaling Laboratory, Sir William Dunn School of Pathology. University of Oxford, South Parks Road, Oxford, OX1 3RE, UK

## Abstract

Src family kinases (SFKs) share highly conserved catalytic domains yet display distinct biological functions, raising the question of how substrate specificity is achieved. Here, we investigate the molecular basis of differential ITAM recognition by Lck and Src, combining cellular assays with structural analysis and docking simulations. In-cell assays demonstrated that, contrary to Lck, Src was completely incapable of phosphorylating the TCR ITAMs when ectopically expressed in a T cell environment. Domain-swapping experiments further revealed that substitution of the Src kinase domain with that of Lck was sufficient to confer ITAM phosphorylation and trigger downstream TCR signaling responses, whereas exchange of adaptor domains had minimal effect. Comparative structural analysis revealed that, despite their overall conserved fold, Lck exhibits a more open and solvent accessible pocket located between the N- and C-lobes of the kinase domain, adjacent to the activation loop, compared to Src. Consistent with this, docking simulations showed that Lck accommodates ITAM peptides in multiple favourable conformations, whereas Src displays a markedly reduced number of non-productive binding poses. Residue-level contact analysis identified a defined interaction surface in Lck, spanning the inter-lobal regions and activation loop. Our results highlight the importance of kinase domain conformational landscape in shaping substrate selectivity and have implications for the rational design of selective SFK inhibitors.

## Introduction

Src family kinases (SFKs) are indispensable transducers of signaling responses, serving a wide variety of receptors and controlling a plethora of functions such as survival, regulation of cell growth, activation, and motility. The best-characterized family members, c-Src, Fyn, Lck and Lyn are amongst the most extensively studied kinase proto-oncogenes and constitute very attractive therapeutic targets due to their central roles in several diseases (1).

Members of the SFK family share a common structural architecture. Starting with a unique domain, the extreme N-terminus (SH4 domain) of which, can undergo lipid modifications, SH3 and SH2 domains with adaptor functions, a regulatory linker sequence followed by the kinase domain, and conclude in a C-terminal inhibitory tail (2).

The enzymatic function of all SFK members is regulated through a highly conserved mechanism, involving specific conformational changes (2). Intramolecular interactions between the SH2 domain and phosphorylated Y527 (amino-acid numbering based on the archetype of SFKs, c-Src), within the C-terminal inhibitory tail, endorse a so-called “closed” conformation, which prevents substrate accessibility to the catalytic pocket. It has been proposed that the closed conformation is further stabilized through the interaction of the SH3 domain with the regulatory linker. Release of these intramolecular blocks by dephosphorylation of Y527 and/or the active displacement of the SH2/SH3 domains by binding interactors forces the kinase into a functional open conformation known as the “primed” form. However, optimal enzymatic activity requires the repositioning of a loop inside the kinase domain, which is achieved via trans-autophsophorylation of Y416. Consequently, the enzymatic activity of SFKs is tuned by a dynamic regulation on the phosphorylation status of the two regulatory tyrosines, Y416 and Y527. Two enzymes with opposing actions have been extensively characterized as the major rheostats of SFK activity. The PTK Csk, responsible for phosphorylating Y527, and the transmembrane PTP CD45 (in haematopoietic cells) (1,2).

SFKs are a highly conserved family with 60-90% sequence identity in their catalytic sites and very similar 3D structures (2). Accordingly, high-throughput screening analyses (3–6) have demonstrated significant, yet not absolute, overlap amongst SFK substrates. Determinants of selectivity have been attributed to tissue-restricted expression and unique patterns of subcellular localization, but importantly, also structural features that influence substrate selection and effective catalysis. To revisit the question of SFK selectivity beyond the conventional *in vitro* settings, we have developed an experimental system that enables monitoring SFK-substrate interactions within a physiological cellular environment.

We selected Lck and c-Src for in-cell substrate specificity comparisons. We reasoned that intrinsic determinants of selectivity would be more clearly resolved by contrasting a T cell–specialized kinase (Lck) with the ubiquitously expressed prototypical SFK c-Src, which is largely absent from resting peripheral T cells (7), than by comparing immune-restricted SFKs such as Lck, Fyn, Lyn, or Hck.

Our data demonstrates that, within an intact cellular environment, Src and Lck can display an astonishing ability for substrate discrimination and that this behaviour is highly dependent on determinants located within their kinase domains. The identification and comprehension of these unique signatures is particularly relevant for the diagnostic and pharmaceutical capitalisation of SFKs, especially within the context of developing potent and clinically safe SFK inhibitors, a task which is met with limited success, primarily due to inadequate inhibitor selectivity towards individual family members (8,9).

## Results

The fundamental importance of Lck in initiating TCR signalling responses is well-established, as is the identification of ITAM tyrosines as its primary and best-characterized substrates (10).

We selected the T-lymphocyte cellular environment where, unlike Src, Lck has been evolutionary selected to operate, to assess selectivity analyses, aspiring to record differences in ITAM-phosphorylating potencies between the two SFKs.

The wild-type sequences of human Lck and Src were introduced in the Lck-deficient Jurkat variant JCaM1.6 via lentiviral gene transfer and using a tetracycline-dependent transcriptional activator. This system of inducible gene expression has been extensively characterized in previous works from our research team ((11,12) and Fig.S1).

Lck- or Src-expressing cells were stimulated with anti-CD3ε for 2min at 37°C and stained with anti-pY142, an antibody specifically recognizing the phosphorylated form of the second tyrosine on the third ITAM of the TCRζ chain (Fig.1B), a well-recognized Lck direct substrate.

To quantify induction of Y142 phosphorylation by the individual kinases, samples were co-stained with a-pY416; this antibody indiscriminately recognizes the phosphorylated form of the activation loop regulatory tyrosines (Y394 in Lck and Y416 in Src). Since the antibody epitope covers a sequence that is absolutely conserved in all SFKs (2), it allows measurements of pY142 within gates of equal levels of SFK activity (gating strategies described in Fig.S1). Collective FACS analysis data (Fig. 1B) revealed that, unlike Lck, Src was completely incapable of inducing Y142 phosphorylation.

**Figure 1.**
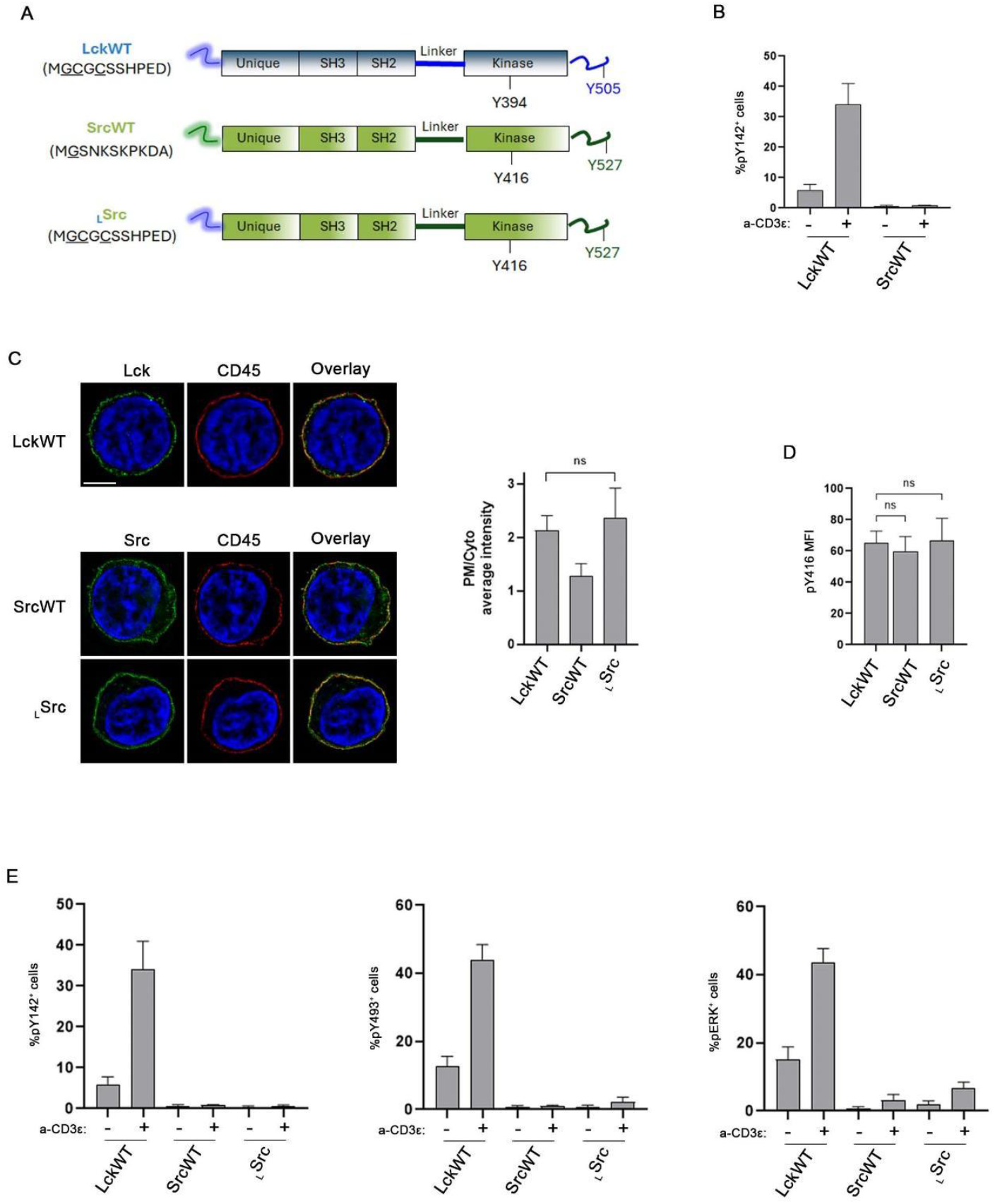
Differential substrate recognition by Lck and Src. **A. Schematic diagrams of Lck and Src domain architecture**. Functional domains and regulatory tyrosines of Lck and Src are color-coded. The SH4 domain amino acid sequences are listed, myristoylation (Glycine) and palmitoylation (Cysteine) sites are underlined. **B. WT Src is unable to induce ITAM phosphorylation in intact T cells**. JCaM1.6 cells expressing Lck or Src wild-type proteins were either left untreated or stimulated with 1μg/ml a-CD3ε (clone UCHT1) for 2min at 37°C and stained with a-pY142 and a-pY416. Graph shows the percentage of cells with levels of pY142 exceeding the -Dox baseline, within pY416+-gated populations (gating strategy described in Fig.S1). Data from four independent experiments. **C. SH4 domain exchange redirects Src subcellular distribution to mimic that of Lck**. SIM images of cells expressing LckWT, SrcWT or _L_Src stained with anti-CD45 (red) and either anti-Lck or anti-Src (green) as indicated. Nuclei were counterstained with DAPI (blue). Adjacent graphs display the PM/Cyto ratio of average intensity fluorescence for each antibody. Number of cells ≥20 from 2 independent experiments. Scale bar: 5μm. **D. SH4 domain exchange does not compromise the levels of active Src**. MFI values of the same cell lines as in B stained with a-pY416 and analysed by FACS (Gating strategy described in Fig.S1). Pooled data from 4 independent experiments. **E. Differential subcellular distribution does not account for the lack of ITAM phosphorylation by Src**. Same cell lines as in B, were either left untreated or stimulated with 1μg/ml a-CD3ε for 2min at 37°C. Samples were stained with a-pY142, a-pY493, (recognizing the phosphorylated form of the activating ZAP-70 tyrosine) a-pERK, (detecting the doubly phosphorylated active status of the ERK1 and ERK2 MAPKs). Graphs display percentage of cells with phosphoantibody fluorescence levels exceeding the -Dox baseline, within pY146+-gated populations. Pooled data from 4 independent experiments. Unpaired Student t test; mean +/-SD; ns: not significant.

The SH4 domain of Lck is a major determinant for its subcellular distribution and lateral residency in discriminating plasma membrane (PM) compartments (12,13). Acyl group modifications within this short 10 amino acid N-terminal sequence, include co-translational myristoylation of Gly2 and the post-translational palmitoylation of Cys3 and Cys5, which appear to be essential for Lck-mediated TCR signaling responses (14,15). Src has been shown to partly reside at the PM, but also, to perinuclear membranes, and in cytoplasm-based vesicles. Similarly to Lck, SrcSH4 is myristoylated on Gly2 but lacks the palmitoylation signals. Instead, it contains a polybasic region involved in membrane association (14,16). Despite their tendency for PM residency, two-colour PALM microscopy of fluorescently-labelled LckSH4 and SrcSH4, showed no significant overlap between these two constructs at the surface of transfected cells, indicating distinct PM compartmentalization.

To exclude that Src inefficiency in promoting Y142 phosphorylation was due to its subcellular distribution, that might prohibit proximity to the TCR complex, we created a chimeric Src construct (_L_Src) in which the Src SH4 domain was substituted by the corresponding sequence of Lck (Fig.1A).

Cells expressing LckWT, SrcWT or _L_Src were analysed by SIM (Fig.1C). CD45 and DAPI staining served as a means of defining ROIs representing the PM and nucleus respectively. Measurements of the average fluorescence, obtained by anti-Lck or anti-Src staining within the defined cellular compartments, were utilized to calculate ratios of PM versus cytoplasmic localization (PM/Cyto) of the specific constructs.

As previously reported, LckWT primarily resided at the plasma membrane, with some anti-Lck fluorescence also being detected intracellularly, whereas SrcWT was proportionally distributed between the PM and cytoplasmic compartments. _L_Src displayed enhanced PM localization compared to SrcWT, with a profile mimicking to that of Lck (Fig.1C).

Displacement of Src from its conventional residence sites did not seem to affect the phosphorylation status of activating tyrosine Y416. Indeed, all three SFK constructs appeared to have comparable levels of their active form, as revealed by quantitative FACs using the anti-pY416 antibody (Fig.1D).

Nevertheless, even when enforced to reside within the same PM compartment as Lck, and thus acquire comparable access to the TCR ITAMs, Src remained unable to phosphorylate Y142. This pattern extended to downstream TCR-mediated signalling events (Fig. 1E), indicating that Src’s inability to phosphorylate ITAM tyrosines is global, rather than restricted to one single tyrosine (Y142). Thus, lack of signal transducing competence towards the TCR machinery is neither due to compromised substrate accessibility, nor inadequate levels of the kinase active form. Rather, it is deduced that intrinsic structural features within Lck define its ability to accommodate the TCR ITAMs for phosphotransfer reactions.

To pinpoint determinants within the Lck structure that would provide substrate specificity, we took advantage of the highly conserved structural architecture of SFKs. Towards this end, Individual domains of Src were sequentially substituted by the corresponding ones of Lck, creating a series of chimeric proteins presented in Fig.2A (detailed description in Table S1). All chimeras were produced on the _L_Src background, to ensure no bias introduced by the divergent subcellular distributions of the two SFKs. To preserve, as adequately as possible, the intramolecular interactions responsible for regulating SFK activity, swaps of the SH3 and SH2 domains, were coupled with corresponding substitutions of the linker and C-terminal inhibitory tail sequences, respectively (Fig.2A and Table S1). Chimeric Src proteins were expressed in the JCaM1.6 line. Importantly, all chimeras exhibited comparable levels of phosphorylation at their regulatory tyrosines (Y416 and Y525: the inhibitory C-terminal tyrosine of Src), indicating that incorporation of the corresponding Lck domains did not disrupt kinase conformation or regulatory integrity (Fig. 2B).

**Figure 2.**
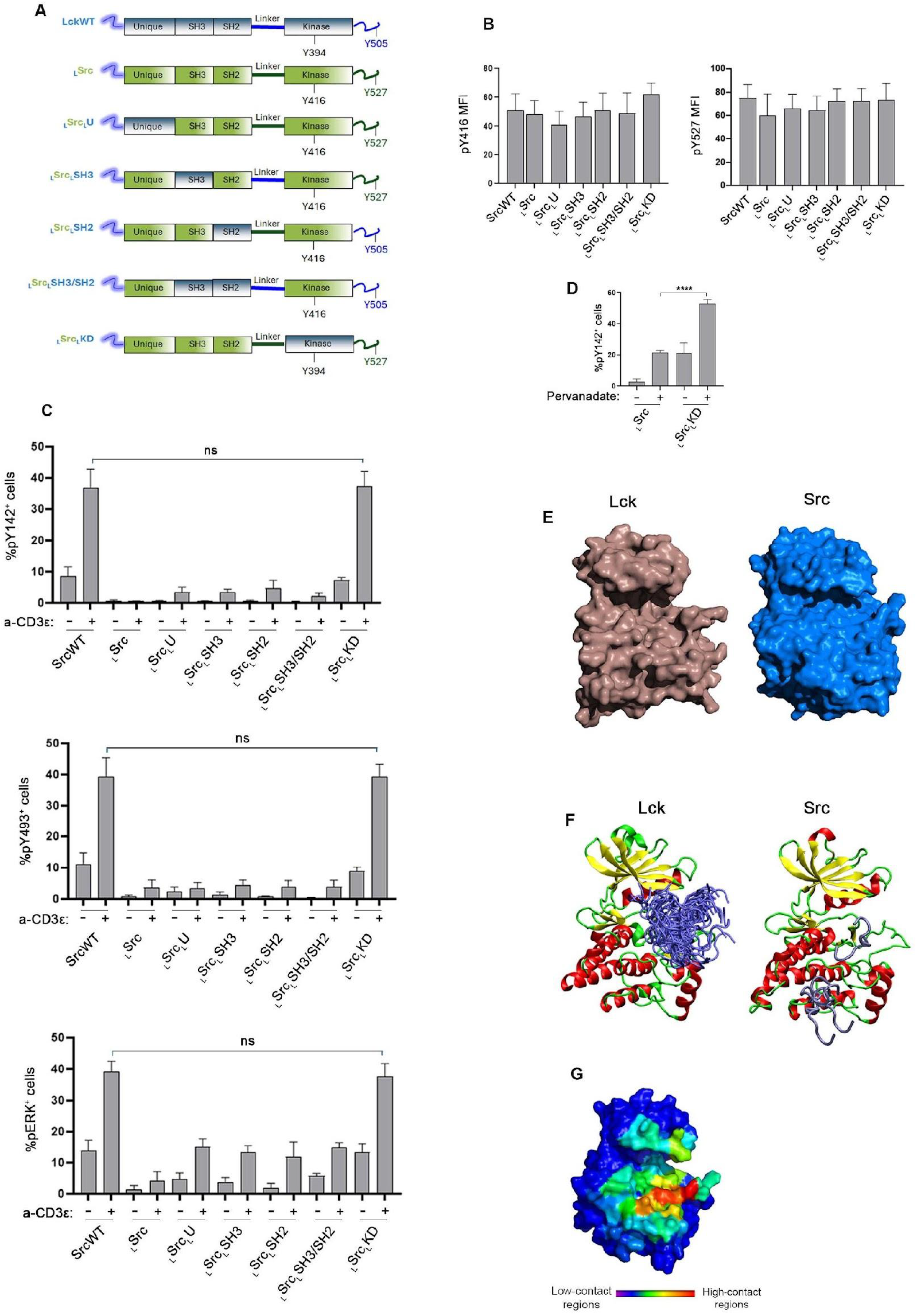
Substrate discrimination features are encoded in the Lck KD. **A. Design of the Src domain-swap chimeras.** Schematic diagrams of Src chimeric proteins. The functional domains and regulatory tyrosines of Lck and Src are designated and colour-coded. **B. Domain swapping did not compromise the conformational integrity of Src.** JCaM1.6 cells expressing SrcWT or chimeric forms were stained with a-pY416 or a-pY527 and analysed by FACS. MFI values of each antibody are displayed in the corresponding graphs as indicated. Pooled data from 4 independent experiments. **C. Incorporation of the Lck kinase domain into Src rescues ITAM phosphorylation and TCR signaling responses**. JCam1.6 cells expressing Lck WT or the indicated Src chimeras were stimulated, stained and analysed as in Fig.1D. Graphs display pooled data from 4 independent experiments. **D. Pervanadate treatment fails to restore the ITAM phosphorylation burden imposed by the Src kinase domain**. Cells expressing _L_Src or _L_Src_L_KD were either left untreated of incubated with 100μM Pervanadate for 10min at 37°C, stained with a-pY142 and a-Src and analysed by FACS. Graph shows the percentage of cells with levels of pY142 exceeding the -Dox baseline, within Src+-gated populations. Data from four independent experiments. **E-G. Intrinsic structural features of the Lck Kinase domain enable optimal ITAM accommodation. E**. Solvent accessible surface X-ray crystal structures of the kinase domains of Lck (light brown) and Src (light blue) showing the presence of a crevice located between the C- and N-lobe of the kinase. **F**. Lck- and Src-ITAM complexes as obtained by docking calculations. The Lck and Src KDs are represented as ribbon and the ITAM peptides as tube (blue marine). **G**. Color-coded surface representations of the Lck-ITAM residue contacts. Two residues are considered in contact when the distance between any of the residue atom pairs is smaller than 6 Å.

The chimeric Src constructs alongside LckWT, were subjected to anti-CD3 stimulation experiments, to assess whether introduction of Lck functional domains would rescue TCR signaling responses (Fig.2C). Incorporation, within the Src protein, of the Lck Unique domain, SH3, SH2 or a combination of the latter two (SH3/SH2), induced somewhat improved yet severely compromised responses and only at very high levels of Src kinase activity (Fig. S2). This pattern changed only when the Src kinase domain (KD) was replaced by the corresponding one of Lck. These data indicate that there are determinants within the Lck kinase domain playing a major part in substrate selectivity towards the ITAMs of the TCR.

To further substantiate the concept that the Src KD is suboptimally configured for accommodating Lck-restricted substrates, we treated the _L_Src and _L_Src_L_KD lines with pervanadate. By inhibiting tyrosine phosphatases, pervanadate drives widespread ITAM and global tyrosine phosphorylation in T cells, effectively bypassing the normal regulatory constraints of the TCR signaling machinery.

Although ITAM phosphorylation was clearly enhanced in _L_Src cells following pervanadate treatment (when compared with a-CD3 stimulations, Fig. 2C), the levels remained substantially low; notably, they only reached baseline phosphorylation levels recorded in untreated _L_Src_L_KD cells (Fig.2D).

To gain insight on the structural basis underlying SFK’s selectivity for substrate recognition, we performed comparative structural analysis and docking simulations. Structural alignment of the crystallographic structures of the kinase domains of Lck (PDB: 3LCK) and Src (PDB: 1YI6) in their active conformations (with phosphorylated activation loop tyrosines Y394 and Y416, respectively), revealed a high degree of similarity (with a root mean square deviation (RMSD) below 1 Å), indicating an overall conserved fold. Nonetheless, notable local differences were observed in the region adjacent to the activation loop, which is implicated in substrate recognition (Fig.2E). Specifically, residues 389–398 of Lck adopt a conformation that is more solvent-exposed compared to the corresponding region in Src. This orientation equips Lck with a wider surface at the interface between the N- and C-lobes of the kinase domain. Pocket detection analysis identified a cavity located in the inter-lobal region in both kinases. However, the pocket volume was substantially larger in Lck compared to Src (9180 Å^3^ and 5736 Å^3^, respectively), consistent with the more solvent-exposed conformation of Lck. These observations suggest that subtle structural differences in the kinase domain may influence substrate accessibility.

To evaluate the functional implication of these structural features, we performed unbiased docking simulations with the aforementioned crystal structures of Lck and Src kinase domains used as receptors, and the NMR-derived structure of an ITAM peptide (PDB: 1CV9), using the HEX server (Fig.2F).

The Lck-ITAM docking simulations produced a large number of favorable poses, with the top 40 solutions shown in Fig.2F, suggesting that the Lck kinase domain can accommodate ITAM peptides in multiple orientations. In contrast, ITAM docking to Src produced only 4 acceptable poses, none of which engaged the putative binding pocket identified from the structural analysis, suggesting that the conformation of Src restricts access of ITAMs to this region.

To further characterize the interaction interface between Lck and the ITAM peptide, residue-residue contacts were calculated for each docking solution (Fig.2G and S3). The surface representation in Fig.2G is color-coded according to the average number of contacts per residue and highlights a prominent interaction region within the pocket between the N- and C-lobes. Consistently, residues 250–257, 278, 326, 329, 364, 366, 385, 397–407, and 437–447 of Lck are most frequently involved in contacts with the ITAM peptide across the ensemble of docking poses. Together, these analyses indicate that the structural architecture of the Lck kinase domain provides a surface that can accommodate the ITAM peptide, whereas the corresponding conformation in Src prohibits accessibility.

## Discussion

This work presents new mechanistic insight into the substrate preference of SFKs. Over the past decades, *in vitro* assays with short synthetic peptides, high-throughput *in vitro* screens using Surface Plasmon Resonance-based profiling of hundreds of substrates, as well as large-scale peptide library screens, combined with next-generation sequencing (3–6,17), have revealed that SFKs share a large pool of common substrates, indicating widespread redundancy and functional overlap. Our data demonstrate that when expressed on their native form within an intact cellular environment, Lck and Src exhibit astonishing selectivity towards the TCR ITAMs and triggering downstream signaling cascades.

Rigorous examination of features within the Src structure, that prohibit ITAM recognition and phosphorylation, excluded potential involvement of its, differential to Lck, subcellular distribution and thus proximity to its substrates.

The adaptor domains of Lck and Src play a critical role in shaping their catalytic output, by mediating and/or stabilizing selective interactions with specific substrates. Notably, the SH2 domain of Lck can additionally interact with anionic PM lipids, thereby providing an additional layer of spatiotemporal regulation for the recruitment of substrates, such as the TCRζ chain and ZAP-70 (18,19). Nonetheless, substitution of the Src SH3, SH2 domains or a combination thereof, with those of Lck, very modestly enhanced Y142 phosphorylation, and this was observed only at the very high levels of Src chimera expression. In fact, only incorporation of the Lck kinase domain, bestowed to Src the ability to phosphorylate the TCR ITAMs and trigger downstream signalling responses, highlighting intrinsic selectivity at the catalytic level.

Structural and docking analyses suggest that structural accessibility and pocket architecture play a critical, and previously unappreciated role in defining specificity towards ITAM motifs. While the global folds of Lck and Src are nearly identical, the distinct conformation of the Lck activation loop creates an expanded and solvent-accessible pocket between the N- and C-lobes. This structural feature enables the accommodation of ITAM peptides in multiple binding orientations, as reflected by the large number of favourable docking poses and the extensive residue-level contact network observed in Lck. In contrast, the more restricted conformation of the Src activation loop limits both the accessibility and adaptability of the corresponding binding region, resulting in a dramatically reduced number of docking solutions that fail to engage the putative substrate-binding pocket.

Our results are broadly consistent with the structural model of the Lck-ITAM complex proposed by Shah et al. (3). This study reports that Lck and Src exhibit subtle sequence preferences particularly in terms of electrostatic determinants; Src favors negatively charged substrates, whereas Lck preferentially phosphorylates substrates enriched in positively charged residues. In both cases, docking simulations of Lck with the ITAM peptide reveal a defined binding mode within the cleft between the N- and C-lobes engaging a surface that extends beyond the catalytic site. However, our simulations reveal a higher degree of conformational plasticity, with multiple favourable binding poses populating the pocket, introducing a structural constraint that helps explaining why ITAM binding is favored in Lck but not in Src, before even considering sequence preferences.

Our comparative analysis of Lck and Src complements current models by showing that specificity emerges from the interplay between electrostatic complementarity and by activation loop-dependent pocket accessibility and conformational flexibility. In this context, Lck appears to be uniquely optimized for ITAM recognition through a combination of favourable amino acid sequence selection (3) and expanded substrate-binding conformation, that enable ITAM accommodation and contact formation. Although our analyses rely on the use of rigid structural models and cannot fully capture the dynamic nature of kinase-substrate interactions, they nevertheless provide a plausible structural framework for interpreting the differential capacity of Lck and Src to accommodate ITAM substrates.

The question of substrate recognition determinants amongst SFKs remains a relevant question, particularly within the context of their therapeutic capitalization. Despite decade-long efforts, the clinical use of SFK inhibitors has been hindered by their poor selectivity. Nevertheless, the field remains active and has been shifting towards rational design strategies guided by structural insights of substrate recognition and kinase activity regulation. Notably, the most selective new compounds, including Src substrate-competitive inhibitors, exploit subtle conformational and structural features that distinguish individual SFKs (8,20). In this context, identifying isozyme-specific structural determinants of substrate selectivity has become a key objective. Our data, highlighting distinct kinase domain properties between Lck and Src, provide additional structural insight that may support next-generation design strategies.

## Materials and Methods

### Antibodies and reagents

The anti-pY416-Src, anti-pY527-Src rabbit pAbs, anti-Src rabbit pAb and anti-P-p44/42 (T202/Y204) MAPK (E10) mouse mAb directly conjugated to AlexaFluor 647 were from Cell Signaling Technology; anti-Lck (3A5) mouse mAb, anti-CD45 (YAML 501.4) rat pAb, were from Santa Cruz Biotechnology; anti-Lck rabbit pAb was from Novus biologicals; anti-v-Src (Ab-1) mouse mAb was from Merck Millipore. Anti-pY142-ζ (K25-407.69) coupled to AlexaFluor 647, from BD Biosciences; anti-CD3ε (UCHT1), used for stimulation experiments, was from Biolegend. Secondary Abs for FACS analysis and SIM were: AlexaFluor 594 donkey anti-rat IgG, AlexaFluor 488 goat anti-rabbit IgG and AlexaFluor 647 goat anti-rabbit IgG from Thermo Fischer Scientific. cDNAs of Src domain-swap chimeras were custom made and purchased from Invitrogen, GeneArt® Strings™ DNA Fragments.

### Generation of Tet-On inducible cell lines

Stable, inducible cell lines were generated using the Lenti-X Tet-On Advanced inducible Expression System (Clonetech) according to the manufacturer’s instructions. cDNAs of human LckWT, SrcWT and chimeric constructs were cloned into the lentiviral vector pLVX-Tight-Puro. The packaging cell line HEK293 was maintained at 37°C in DMEM complete media supplemented with10% FCS and was used for the production of recombinant lentiviruses. Specifically, 80% confluent HEK293 were transfected with a mixture of the lentiviral packaging plasmids pSVG and pSPAX2 and either pLVX-Tight-Puro containing the Lck constructs or the PLVX-Tet-On-Advanced vector driving the expression of the Tet repressor protein. 48h after transfection, the supernatants containing lentiviral particles were collected and the two batches of lentiviruses were combined and used to transduce the Lck-deficient JCaM1.6 parental cells. 48h post-infection the cells were put under selection by Puromycin (10μg/ml) and geneticin (1mg/ml) and maintained in culture at 37°C in RPMI supplemented with 10% FCS. Expression of the Lck constructs was induced by 1μg/ml Doxycycline (Sigma-Aldrich) added to the cell culture medium, routinely 24h prior to each experiment.

### Cells, induction of Lck expression, stimulation and Flow Cytometry

Cell lines were cultured at a density of 300,000 cells/ml in RPMI-1640-10% FCS. Induction of protein expression was initiated by the addition of 1μg/ml Doxycycline (SIGMA) to the culture media, 24h prior to each experiment. Typically, for anti-CD3ε stimulation, cells were incubated with 1μg UCHT1 for 2min at 37°C. Single-cell suspensions were fixed for 10min with BD Phosflow™ Fix Buffer I (BD Biosciences) prior to 10min permeabilization with permeabilization buffer (0.5% BSA, 0.5% OmniPur Triton X-100 Surfactant (Millipore) in PBS). Primary antibodies, diluted in permeabilization buffer, were added to the cells for 1h, followed by three washes in permeabilization buffer and the addition of the corresponding secondary antibodies (again diluted in permeabilization buffer). After a final series of washes cells were analysed on a on a BD Accuri™ C6 Plus Flow Cytometer. Acquired data were analysed by the FlowJo Software (BD Biosciences). Statistical analysis was performed with Prism (GraphPad Software).

### Immunostaining, 3D-SIM, image acquisition and analysis

Single-cell suspensions were immobilized on poly-L-Lysine (Sigma-Aldrich)-coated high precision glass coverslips (Marienfeld) for 10min in a cell culture incubator. Cells were fixed for 10min with 4% Paraformaldehyde (PFA) and permeabilized with PBS/0.5% Triton-X100 for 5min. After blocking with PBS/1%BSA for 15 min, cells were stained for 1h at room temperature (RT) with the indicated primary Abs. The corresponding fluorochrome-conjugated secondary Abs were added for 1h. Nuclei were counterstained with 1 μg/ml DAPI (Sigma-Aldrich), and coverslips were mounted to microscopy slides with ProLong Gold anti fade reagent (Vectashield).

SIM was performed on an OMX V3 BLAZE microscope (GE Healthcare) using 405-, 488- and 592-nm solid-state multimode lasers and a 60x/1.42 oil UPlanSApo objective (Olympus). Multi-channel images were captured sequentially by Photometrics Cascade back-illuminated EMCCD cameras (Photometrics).1μm SIM stacks were acquired at 125nm z-distance, with 15 raw images per plane (three angles, five phases) resulting in 120 raw images in total, for each sample.

Calibration measurements of 0.2μm diameter Tetraspec fluorescent beads (Life Technologies) were used to obtain alignment parameters subsequently utilized to align images from the different color channels (OMXEditor software). A total of 120 raw image data per sample were computationally reconstructed using the SoftWoRx 6.0 software package (Applied Precision) to obtain super-resolution image stacks with a resolution of 100nm in x and y and approximately 120nm in z.

Images were converted to 16-bit average intensity composites and defined regions of interest (ROI) were segmented using binary image masks drawn according to the fluorescence obtained from the DAPI and anti-CD45 staining. The CD45 staining mask defined the PM ROI whereas and the DAPI staining mask designated the nuclear ROI. The area between the PM and nuclear masked regions was defined as the cytoplasmic ROI. Measurements of the average fluorescence intensities within the respective PM and cytoplasmic ROIs were used to calculate the plasma membrane/cytoplasmic ratios for the staining of anti-Lck, anti-Src, anti-pY416 and anti-pY505 antibodies. All image processing was performed using in-house ImageJ scripts.

### Structural analysis and docking simulations

Structural alignment of the kinase domains of Lck and Src was performed using the crystal structures of their active conformations (PDB IDs:3LCK and 1YI6, respectively). The structures were superimposed using the alignment function of PyMOL (Schrödinger, LLC), and the root mean square deviation (RMSD) between the catalytic cores was calculated to assess overall structural similarity.

Potential ligand-binding cavities in the kinase domains were identified using the Fpocket software package . Pocket volumes for Lck and Src were compared to assess differences in accessibility of the substrate-binding region. Solvent-accessible surface representations were generated by PyMOL to visualize the spatial relationship between the activation loop and the inter-lobal cleft of the kinase domains.

Docking calculations were performed using the HEX docking server. The kinase domains of human Lck (PDB ID:3LCK) and Src (PBD ID: 1YI6) were used as receptors, while the ITAM peptide structure was obtained from the NMR structure PBD ID: 1CV9. Docking calculations were performed without restraints, using default parameters, including shape and electrostatic correlations with a search order parameter at 25. The resulting docking poses were ranked according to the scoring function implemented in HEX. The highest-ranking docking solutions were analyzed to identify productive ligand-kinase domain interactions.

To characterize the interaction interface between Lck and the ITAM peptide, residue-residue contacts were calculated for the ensemble of docking solutions. Two residues were considered to be in contact when the distance between any pair of atoms was less than 6 Å. Contact frequencies were calculated for each Lck residue across the ensemble of docking poses, allowing identification of regions most frequently involved in ITAM peptide interactions.

## References

*1) HexServer: an FFT-based protein docking server powered by graphics processors. G. Macindoe, L. Mavridis, V. Venkatraman, M.-D. Devignes, D.W. Ritchie (2010). Nucleic Acids Research, 38, W445-W449*

*2) Fpocket: An open-source platform for ligand pocket detection, V. Le Guilloux, P. Schmidtke and P. Tuffery, BMC Bioinformatics, 2009, 10:168*

*3) The PyMOL Molecular Graphics System, Version 1.7.4 Schrödinger, LLC.*

## References

1. Eshaq AM, Flanagan TW, Hassan S-Y, Al Asheikh SA, Al-Amoudi WA, Santourlidis S, Hassan S-L, Alamodi MO, Bendhack ML, Alamodi MO, et al. Non-Receptor Tyrosine Kinases: Their Structure and Mechanistic Role in Tumor Progression and Resistance. Cancers (Basel) (2024) 16:2754. doi: 10.3390/cancers16152754

2. Boggon TJ, Eck MJ. Structure and regulation of Src family kinases. Oncogene (2004) 23:7918–7927. doi: 10.1038/sj.onc.1208081

3. Shah NH, Löbel M, Weiss A, Kuriyan J. Fine-tuning of substrate preferences of the Src-family kinase Lck revealed through a high-throughput specificity screen. Elife (2018) 7: doi: 10.7554/eLife.35190

4. Deng Y, Alicea-Velázquez NL, Bannwarth L, Lehtonen SI, Boggon TJ, Cheng H-C, Hytönen VP, Turk BE. Global Analysis of Human Nonreceptor Tyrosine Kinase Specificity Using High-Density Peptide Microarrays. J Proteome Res (2014) 13:4339–4346. doi: 10.1021/pr500503q

5. Li A, Voleti R, Lee M, Gagoski D, Shah NH. High-throughput profiling of sequence recognition by tyrosine kinases and SH2 domains using bacterial peptide display. Elife (2023) 12: doi: 10.7554/eLife.82345

6. Takeda H, Kawamura Y, Miura A, Mori M, Wakamatsu A, Yamamoto J, Isogai T, Matsumoto M, Nakayama KI, Natsume T, et al. Comparative Analysis of Human Src-Family Kinase Substrate Specificity in Vitro. J Proteome Res (2010) 9:5982–5993. doi: 10.1021/pr100773t

7. Branch DR, Mills GB. pp60c-src expression is induced by activation of normal human T lymphocytes. J Immunol (1995) 154:3678–85.

8. Su Y, Zhu K, Wang J, Liu B, Chang Y, Chang D, You Y. Advancing Src kinase inhibition: From structural design to therapeutic innovation - A comprehensive review. Eur J Med Chem (2025) 287:117369. doi: 10.1016/j.ejmech.2025.117369

9. Aleshin A, Finn RS. SRC: A Century of Science Brought to the Clinic. Neoplasia (2010) 12:599–607. doi: 10.1593/neo.10328

10. Woessner NM, Uleri V, Stepanek O, Minguet S. The TCR and LCK: foundations for T-cell activation and therapeutic innovation. Front Immunol (2026) 16: doi: 10.3389/fimmu.2025.1737013

11. Koutras N, Morfos V, Konnaris K, Kouvela A, Shaukat A-N, Stathopoulos C, Stamatopoulou V, Nika K. Integrated signaling and transcriptome analysis reveals Src family kinase individualities and novel pathways controlled by their constitutive activity. Front Immunol (2023) 14: doi: 10.3389/fimmu.2023.1224520

12. Porciello N, Cipria D, Masi G, Lanz A-L, Milanetti E, Grottesi A, Howie D, Cobbold SP, Schermelleh L, He H-T, et al. Role of the membrane anchor in the regulation of Lck activity. Journal of Biological Chemistry (2022) 298:102663. doi: 10.1016/j.jbc.2022.102663

13. Soares H, Henriques R, Sachse M, Ventimiglia L, Alonso MA, Zimmer C, Thoulouze M-I, Alcover A. Regulated vesicle fusion generates signaling nanoterritories that control T cell activation at the immunological synapse. Journal of Experimental Medicine (2013) 210:2415–2433. doi: 10.1084/jem.20130150

14. Resh MD. Myristylation and palmitylation of Src family members: The fats of the matter. Cell (1994) 76:411–413. doi: 10.1016/0092-8674(94)90104-X

15. Kabouridis PS. S-acylation of LCK protein tyrosine kinase is essential for its signalling function in T lymphocytes. EMBO J (1997) 16:4983–4998. doi: 10.1093/emboj/16.16.4983

16. Redmond T, Brott BK, Jove R, Welsh MJ. Localization of the viral and cellular Src kinases to perinuclear vesicles in fibroblasts. Cell Growth Differ (1992) 3:567–76.

17. Sicilia RJ, Hibbs ML, Bello PA, Bjorge JD, Fujita DJ, Stanley IJ, Dunn AR, Cheng H-C. Common in Vitro Substrate Specificity and Differential Src Homology 2 Domain Accessibility Displayed by Two Members of the Src Family of Protein-tyrosine Kinases, c-Src and Hck. Journal of Biological Chemistry (1998) 273:16756–16763. doi: 10.1074/jbc.273.27.16756

18. Gao M, Skolnick J. Predicting protein interactions of the kinase Lck critical to T cell modulation. Structure (2024) 32:2168-2179.e2. doi: 10.1016/j.str.2024.09.010

19. Sheng R, Jung D-J, Silkov A, Kim H, Singaram I, Wang Z-G, Xin Y, Kim E, Park M-J, Thiagarajan-Rosenkranz P, et al. Lipids Regulate Lck Protein Activity through Their Interactions with the Lck Src Homology 2 Domain. Journal of Biological Chemistry (2016) 291:17639–17650. doi: 10.1074/jbc.M116.720284

20. Breen ME, Steffey ME, Lachacz EJ, Kwarcinski FE, Fox CC, Soellner MB. Substrate Activity Screening with Kinases: Discovery of Small-Molecule Substrate-Competitive c-Src Inhibitors. Angew Chem Int Ed (2014) 53:7010–7013. doi: 10.1002/anie.201311096

