## Supplementary Material for "Molecular determinants of differential substrate selection between the Src family kinases Lck and Src"

### Supplemental Information

Fig.S1

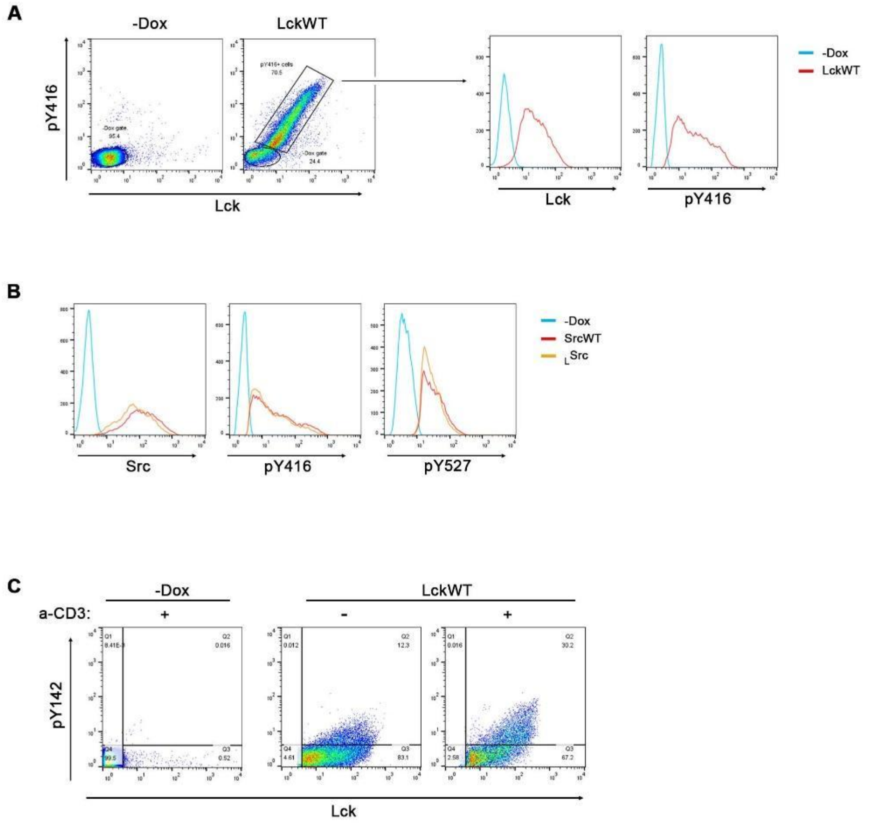


**Figure S1. Gating strategy and antibody validation.**

**A**. JCaM1.6 cells transduced with the LckWT construct were cultured for 48h in the absence or presence of Dox, stained with a-Lck and a-pY416 and analysed by FACS. Single cells (from FSC-A versus FGC-H dot plots-not shown) positive for Lck/pY416 expression were selected based on gating set by respective antibodies' fluorescence in the -Dox samples. Graphs presented in the manuscript, display MFI values or frequencies of cells within the pY416$^{+}$ subsets (or pY527$^{+}$ cells in the corresponding experiments). This gating strategy was followed in all FACS-based analyses throughout the study. Adjacent overlay histograms show the expression levels of Lck, and its active form as indicated.

**B**. Overlay histograms of cells expressing SrcWT, $_{L}$Src or a mix of their -Dox counterparts stained with a-Src, a-pY416 and a-pY527

**C**. Gating strategy for stimulation experiments. LckWT-expressing cells and their -Dox counterparts were incubated in the presence or absence of 1μg/μl anti-CD3ε (clone UCHT1) for 2min at 37°C. Samples were stained with a-pY142 and a-pY416 and analysed by FACS. pY416+ populations (isolated as in A) were further gated based on baseline fluorescence of the indicated antibodies in the -Dox sample. Percentage of cells in the upper right quadrant (Q2) represent the pY142$^{+}$ populations.

Fig.S2

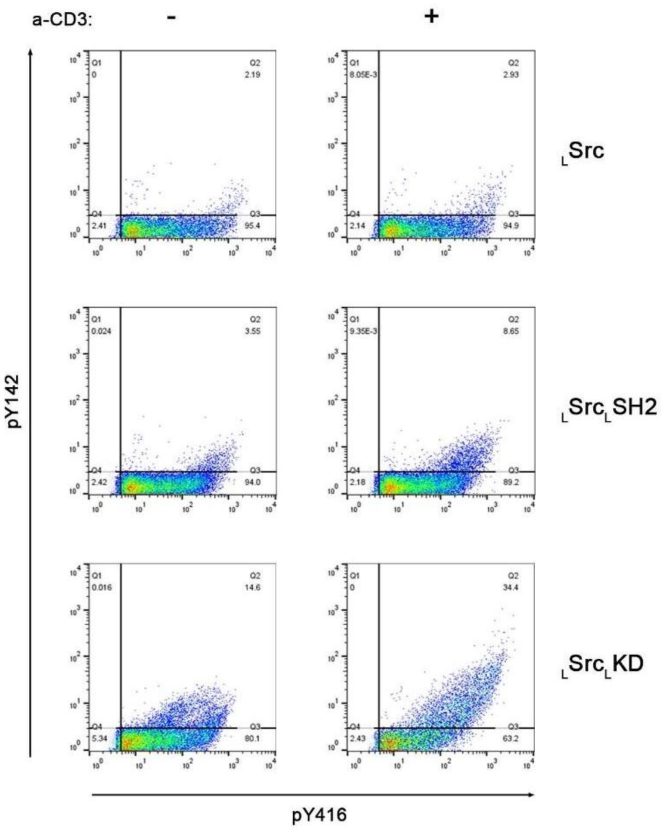


**Figure S2**. Representative 2D FACS plots of JCAM1.6 lines expressing the indicated constructs, stimulated with 1μg/ml a-CD3ε (clone UCHT1) for 2min at 37°C and stained with a-pY142 and a-pY416. Gating as in Fig.S1. Only $_{L}$Src$_{L}$SH2 is displayed as a representative profile for the swap chimeras bestowing a limited improvement on pY142 only at very high levels of SFK activity, as the rest of the constructs exhibit very similar patterns.

Fig.S3

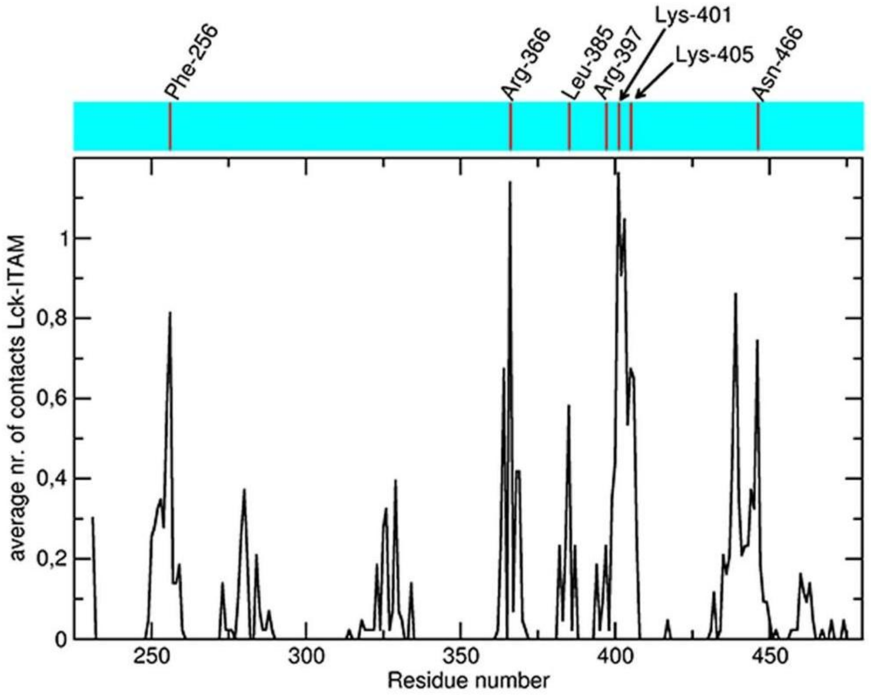


**Figure S3.** Average number of residue-residue contacts for all 40 ITAM poses with respect to any Lck residue, as a function of the Lck residue number. Peaks correspond to high Lck-ITAM contact regions. Overlaid are the corresponding ITAM-Lyn interactions as reported by Gaul and Colleagues (1). Numbering on top is from the Lyn sequence.

*Reference*

*1. Gaul BS, Harrison ML, Geahlen RL, Burton RA, Post CB. Substrate recognition by the Lyn protein-tyrosine kinase. NMR structure of the immunoreceptor tyrosine-based activation motif signaling region of the B cell antigen receptor. J Biol Chem. 2000 May 26;275(21):16174-82. doi: 10.1074/jbc.M909044199. PMID: 10748115.*

**Table S1**. Src domain-swap chimeras[1]

| **Chimeric Protein** | **Fusion site between Src and Lck** | |
|---|---|---|
| | **N-terminal** | **C-terminal** |
| <u>$_{L}$Src$_{L}$U</u> | **GCGCSSHPEDDWME...** | ...***GSNPPAS***PLAGGVTTF... |
| <u>$_{L}$Src$_{L}$SH3</u> | ...TSPQRAGPL***QDNLVIA*** | ***...NFVAKAN***SIQAEEWY... |
| **(Lck Linker)** | ...CHRLTTVC***QTQKPQK***... | ...***KPWWEDE***WEIPRES... |
| <u>$_{L}$Src$_{L}$SH2</u> | ...NYVAPSDS***LEPEPWF***... | ...***CTRLSRP***CPTSKPQT... |
| **(Lck C-terminal tail)** | ...YLQAFLED***FFTATEGQYQPQP*** | |
| <u>$_{L}$Src$_{L}$SH3/SH2</u> | ...TSPQRAGPL***QDNLVIA***... | ...***KPWWEDE***WEIPRES... |
| **(Lck Linker)** | ...CHRLTTVC***QTQKPQK***... | ...***KPWWEDE***WEIPRES... |
| **(Lck C-terminal tail)** | ...YLQAFLED***FFTATEGQYQPQP*** | |
| $_{L}$Src$_{L}$KD | ...GLAKDAWE***VPRETLKL***... | ...***YLRSVLE***DYFTSTEP... |

[1]In all chimeras lack the first 16 amino acids of human Src have been replaced by the SH4 domain of Lck. **Bold characters**: sequence of the Lck region or domain according to UniProtKB, Regular characters: amino acid sequence of Src. Conserved amino acids between Src and Lck at the borders of the fusion site of the topological domains are designated in red.
